# Real-time measurement of E2:ERα transcriptional activity in living cells

**DOI:** 10.1101/844761

**Authors:** Manuela Cipolletti, Stefano Leone, Stefania Bartoloni, Claudia Busonero, Filippo Acconcia

## Abstract

Kinetic analyses of diverse physiological processes have the potential to unveil new aspects of the molecular regulation of cell biology at temporal levels. 17β-estradiol (E2) regulates diverse physiological effects by binding to the estrogen receptor α (ERα), which primarily works as a transcription factor. Although many molecular details of the modulation of ERα transcriptional activity have been discovered including the impact of receptor plasma membrane localization and its relative E2-evoked signalling, the knowledge of real-time ERα transcriptional dynamics in living cells is lacking. Here, we report the generation of MCF-7 and HeLa cells stably expressing a modified luciferase under the control of an E2-sensitive promoter, which activity can be continuously monitored in living cells and show that E2 induces a linear increase in ERα transcriptional activity. Ligand-independent (*e.g*., epidermal growth factor) receptor activation was also detected in a time-dependent manner. Kinetic profiles of ERα transcriptional activity measured in the presence of both receptor antagonists and inhibitors of ERα plasma membrane localization reveals a biphasic dynamic of receptor behaviour underlying novel aspects of receptor-regulated transcriptional effects. Finally, analysis of the rate of the dose-dependent E2 induction of ERα transcriptional activity demonstrates that low doses of E2 induce an effect identical to that determined by high concentrations of E2 as a function of the duration of hormone administration. Overall, we present the characterization of sensitive stable cell lines where to study the kinetic of E2 transcriptional signaling and to identify new aspects of ERα function in different physiological or pathophysiological conditions.

## Introduction

The sex hormone 17β-estradiol (E2) is a critical regulator of cell physiology as it controls the homeostasis of female and male reproductive and non-reproductive tissues and organs. The pleiotropic E2 actions depend on the activation of E2 signaling. E2 signaling is triggered by the activation the estrogen receptor α (ERα), which works as a ligand-induced transcription factor (Busonero et al., 2019).

Indeed, in the nucleus, the E2:ERα complex activates the transcription of the genes containing the estrogen response element (ERE) sequence in their promoters. Moreover, E2 can regulate the expression of non-ERE sequence containing genes by inducing the ERα interaction with other transcription factors (Ascenzi et al., 2006). In addition, the regulation of ERα transcriptional activity can be further triggered by other hormones (*e.g*., epidermal growth factor – EGF) in the absence of E2 (Ascenzi et al., 2006). Activation of E2:ERα complex gene transcription requires receptor phosphorylation on the serine (S) residue 118 (Lannigan, 2003; Le Romancer et al., 2011). The S118 ERα phosphorylation is a result of the E2-dependent activation of kinase cascades (*e.g*., PI3K/AKT; ERK/MAPK), which are triggered by the plasma membrane localized ERα (Acconcia et al., 2005a; La Rosa et al., 2012; Pedram et al., 2007). ERα plasma membrane localization occurs because the receptor is palmitoylated on the cysteine (C) residue 447 by specific palmitoyl-acyl-transferases (PAT) (Acconcia et al., 2005a; Adlanmerini et al., 2014; La Rosa et al., 2012; Pedram et al., 2012; Pedram et al., 2007; Sosa et al., 2019). The ERα plasma membrane localization is a pre-requisite for E2-induced ERα transcriptional activity (La Rosa et al., 2012).

In the last years, diverse kinetic analyses have been performed in living cells to monitor real-time cellular responses induced by different extracellular stimuli. Measurement of growth rates of breast cancer cells treated with different hormones (*e.g*., E2, progestins, androgens and corticosteroids) or chemicals (*e.g*., environmental pollutants) revealed complex kinetic proliferation profiles that suggested novel mechanisms of action for each compound (Rotroff et al., 2013). Another method, which has been developed to study protein turnover, allowed to identify in a quantitative time-dependent manner new kinetic mechanistic events required for protein degradation induced by proteolysis targeting chimeras (PROTACs) (Riching et al., 2018).

Therefore, real-time live-cell assays for measuring different parameters of cellular biology holds the potential for the identification of unrecognized biological phenomena underlying the physiological effects of extracellular stimuli (*e.g*., hormones).

Remarkably, to our knowledge, the real-time kinetic evaluation of E2-induced ERα transcriptional activity has never been reported. In turn, we decided to generate a cell line-based model system to detect the ERα transcriptional activity in living cells. To this purpose, we took advantage of the available technology for which the activity of a modified luciferase (nanoluciferase-PEST - NLuc) can be continuously detected in cells loaded with a non-toxic cell-permeable substrate (Hall et al., 2012) and developed MCF-7 and HeLa cell lines stably expressing an E2-responsive NLuc reporter gene construct.

Here, we report the characterization of ERα transcriptional activity in live-cells by different hormones (*i.e*., E2 and EGF) as well as the real-time kinetic analysis of the impact of ERα plasma membrane localization on the E2-induced ERα transcriptional activity.

## Materials and Methods

### Cell culture and reagents

17β-estradiol (E2), epidermal growth factor (EGF), 4OH-tamoxifen (Tam), DMEM (with and without phenol red), fetal calf serum, charcoal stripped fetal calf serum (DCC) and the palmitoyl-acyl-transferase (PAT) inhibitor 2-bromohexadecanoic acid (2-bromo-palmitate; 2-Br) [IC_50_ of ∼4 µM] (Varner et al., 2003) were purchased from Sigma-Aldrich (St. Louis, MO). Bradford protein assay kit as well as anti-mouse and anti-rabbit secondary antibodies were obtained from Bio-Rad (Hercules, CA). Antibodies against ERα (F-10 mouse – for WB), pS2 (FL-84 rabbit) and cathepsin D (H75 rabbit) were obtained from Santa Cruz Biotechnology (Santa Cruz, CA); anti-vinculin antibody was from Sigma-Aldrich (St. Louis, MO). All other antibodies were purchased by Cell Signalling Technology Inc. (Beverly, MA, USA). Chemiluminescence reagent for Western blotting was obtained from BioRad Laboratories (Hercules, CA, USA). Nano-Glo® Endurazine™ was purchased from Promega (Promega, Madison, MA, USA). All the other products were from Sigma-Aldrich. Analytical- or reagent-grade products were used without further purification. The identities of all the used cell lines [*i.e*., human breast carcinoma cells (MCF-7 and T47D-1) and human cervix carcinoma cells (HeLa)] were verified by STR analysis (BMR Genomics, Italy).

### Plasmids and Cloning

The reporter plasmid 3xERE TATA, the pcDNA flag 3.1 C, the pcDNA flag-ERα, the pcDNA flag-ERα C447A, and the pcDNA flag-ERα S118A were previously described (La Rosa et al., 2012). In order to generate the reporter plasmid pGL2Basic Neo_NLucPest_3xERE TATA containing the Nanoluciferase-PEST (NLuc) gene under the control of the 3xERE TATA promoter, the NLuc gene was first excised by the pNL2.2 (Promega, Madison, MA, USA) using KpnI/BamHI sites and cloned into the pGL2Basic Neo 3xERE TATA (a generous gift of Dr Wilson) (Wilson et al., 2004) using KpnI/BamHI sites. The resulting plasmid was the pGL2Basic Neo_NLucPest. Next, the fragment containing the 3xERE TATA was excised by the pGL2Basic Neo 3xERE TATA cut KpnI/HindIII and cloned into the pGL2Basic Neo_NLucPest plasmid cut KpnI/HindIII to finally generate the pGL2Basic Neo_NLucPest_3xERE TATA (Fig. 1).

**Figure 1.**
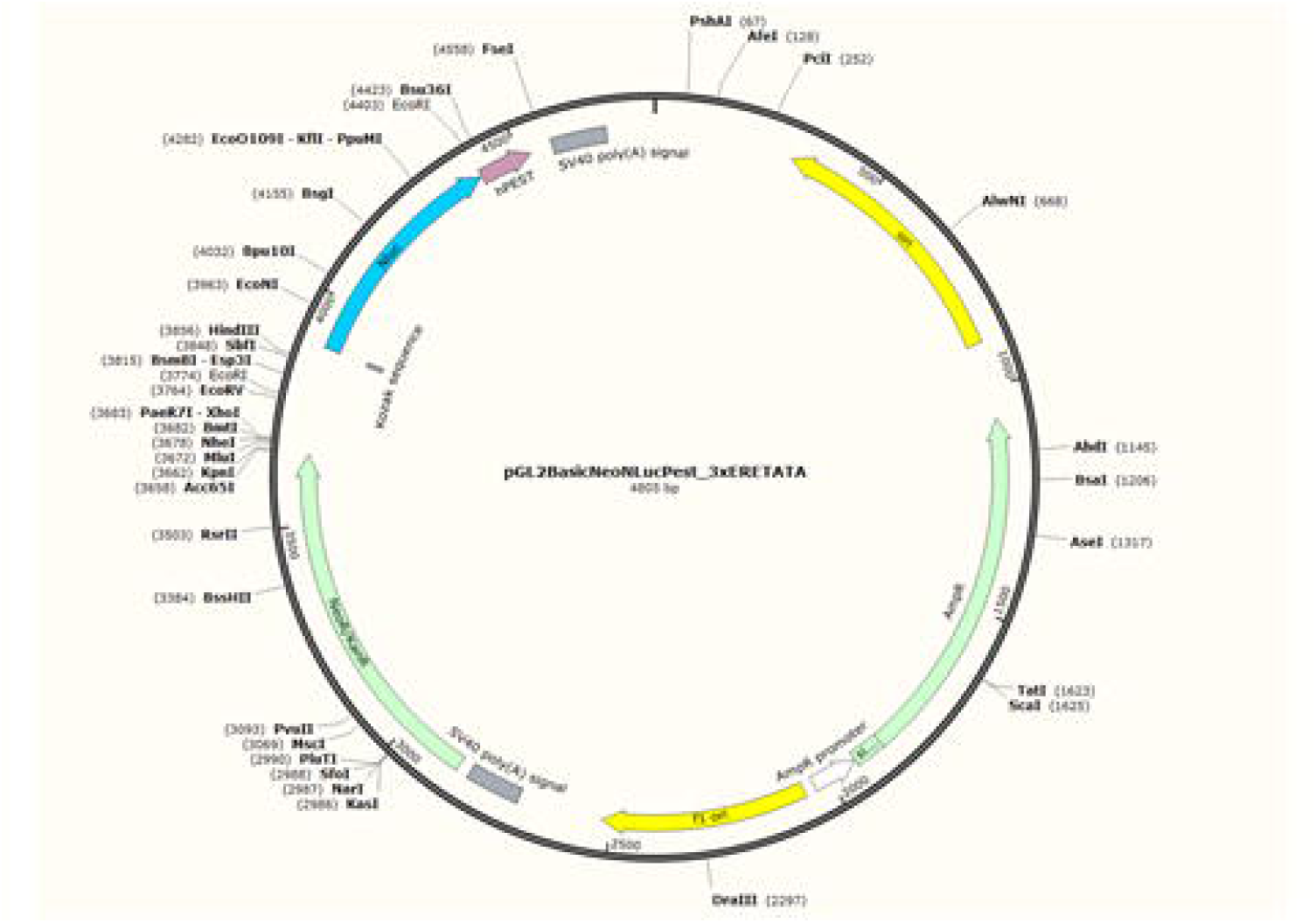
The pGL2Basic Neo_NLucPest_3xERE TATA plasmid. Schematic of the produced plasmid map used to generate the MCF-7 and HeLa ERE-NLuc stable cell lines. Important genetic determinants and restriction enzymes are indicated.

### Generation of stable MCF-7 and Hela ERE-NLuc cell lines

MCF-7 and HeLa cells were transfected with pGL2Basic Neo_NLucPest_3xERE TATA using lipofectamine 2000 (Thermofisher) reagent according to the manufacturer’s instructions. Twenty-four hours after transfection medium was changed and the selection antibiotic was added. In particular, MCF-7 ERE-NLuc cells were generated by using G418 (500 μg/ml) while HeLa ERE-NLuc cells were generated by using G418 (750 μg/ml). Pooled cloned were used for the experiments. Selection antibiotic was left in growing medium while each experiment was performed in the absence of antibiotic.

### Growth Curves

The xCELLigence DP system ACEA Biosciences, Inc. (San Diego, CA) Multi-E-Plate station was used to measure the time-dependent response to E2 by real-time cell analysis (RTCA). Each experimental condition was tested in quadruplicate. A detailed description of the instrument and the relative software has been previously published (Rotroff et al., 2013). Briefly, the instrument measures the electric impedance of the cells on the well surface. The software transforms the measured value of the electric impedance in an a-dimensional parameter called Cell Index (C.I.). Increased electric impedance and consequently and increased C.I. is proportional to an increase in the number of cells. C.I. normalized for each well at time 0 (*i.e*., normalized C.I.) is the parameter used to follow cell proliferation, according to the software manufacturer’s instruction. MCF-7, T47D-1 and MCF-7 ERE-NLuc cells were seeded in E-Plates 96 in growing medium. After overnight monitoring of growth once every 15 min, medium was changed, and cells were grown in 1% DCC medium in the presence or in the absence of E2 (1 nM) and remained in the medium until the end of the experiment. Cellular responses were then recorded once every 15 min for a total time of 72 hours.

For comparison of the effect of E2 in each cell line, the ratio between the normalized cell index (NCI) (obtained by the ACEA Biosciences software) of the mean value for the E2-treated samples and the NCI of the mean value for control sample was calculated and shown as a function of time.

### Real-time measurement of NanoLucPest expression

MCF-7 ERE-NLuc cells were seeded in 96 well plates (5000 cells/well). Twenty-four hours after plating, medium was changed, and cells were grown in 1% DCC medium for 24 hours and then stimulated with E2. Each experimental condition was plated in triplicate and 3 wells were always treated with fulvestrant (ICI182,240) (Sigma Aldrich) in order to measure the basal ERα transcriptional activity. HeLa ERE-NLuc cells were transfected with the indicated ERα encoding plasmids or with vector control. Twenty-four hours after transfection cells were seeded in 96 well plates (5000 cells/well) and subsequently treated as described for MCF-7 ERE-NLuc cells. Nano-Glo® Endurazine™ was added according to manufacturer’s instruction in 50µl as final experimental volume together with ligand and/or inhibitor administration. Plates were then transferred into a Tecan Spark microplate reader (Switzerland) set to 37°C and 5% CO_2_. Light emission (released light units - RLU) was measured for 24 hours every other 5 minutes. For calculations, each data point was subtracted of the RLU mean value of the 3 fulvestrant treated samples at each time point. Next each value was subtracted with the value of the corresponding sample at time 0. The mean value of each experimental condition was calculated and subsequently the ratio between each experimental condition and the control condition was obtained. Each experiment was done twice in duplicate.

### Western Blotting Assays

Before any cellular and biochemical assay, cells were grown in 1% DCC medium for 24 hours and then stimulated with E2 at the indicated time points and doses. Cells were lysed in YY buffer (50 mM HEPES at pH 7.5, 10% glycerol, 150 mM NaCl, 1% Triton X-100, 1 mM EDTA, 1 mM EGTA) plus protease and phosphatase inhibitors. Western blotting analyses were performed by loading 20-30 µg of protein on SDS-gels. Gels were run and transferred to nitrocellulose membranes with Biorad Turbo-Blot semidry transfer apparatus. Immunoblotting was carried out by incubating membranes with 5% milk (60 min), followed by incubation o.n. with the indicated antibodies. Secondary antibody incubation was continued for an additional 60 min. Bands were detected using a Biorad Chemidoc apparatus.

### BrdU Incorporation

Bromodeoxiuridine (BrdU) was added in the last 30 minutes to the medium and then cells were fixed, permeabilized, and the histones were dissociated with 2 M HCl as previously described (Darzynkiewicz and Juan, 2001). BrdU positive cells were detected with an anti-BrdU primary antibody diluted 1:100 (DAKO Cytomatation) and with an anti-mouse-Alexa488 conjugated diluted 1:100 (Thermofisher). Both antibodies were incubated for 1 hour at R.T. in the dark. BrdU fluorescence was measured using a CytoFlex flow cytometer and the cell cycle analysis was performed by CytExpert v1.2 software (Beckman Coulter). All samples were counterstained with propidium iodide (PI) for DNA/BrdU biparametric analysis.

### Cell cycle analysis

After treatments, cells were harvested with trypsin, and counted to obtain 10^6^ cells per condition. Then, the cells were centrifuged at 1500 rpm for 5 min at 4°C, fixed with 1 ml ice-cold 70% ethanol and subsequently stained with PI buffer (500 µg/ml Propidium Iodide, 320µg/ml RNaseA, in 0.1% Triton X in PBS). DNA fluorescence was measured using a CytoFlex flow cytometer and the cell cycle analysis was performed by CytExpert v1.2 software (Beckman Coulter).

### Statistical analysis

A statistical analysis was performed using the ANOVA (One-way analysis of variance and Tukey’s as post-test) test with the InStat version 3 software system (GraphPad Software Inc., San Diego, CA). Densitometric analyses were performed using the freeware software Image J by quantifying the band intensity of the protein of interest respect to the relative loading control band (*i.e*., vinculin) intensity. Numerosity of the experiments is given in figure texts. Data are the mean ± standard deviation. In all analyses, *p* values < 0.01 were considered significant but for Western blotting experiments for which *p* values < 0.05 were considered significant.

## Results

### Characterization of MCF-7 NLuc cells

Characterization of the generated MCF-7 ERE-NLuc cells was performed by evaluating the ability of E2 to induce cell proliferation (Castoria et al., 2001), S118 ERα phosphorylation (Ali et al., 1993), ERα degradation (Leclercq et al., 2006) as well as the accumulation of two well-known ERE-containing genes (*i.e*., presenilin 2 – pS2/TFF and cathepsin D – CatD) (Sun et al., 2005).

Growth curves analyses indicated that E2 induced a persistent time-dependent increase in cell number in MCF-7 ERE-NLuc cells (Fig. 2A-green line). Interestingly, E2 was also able to increase the number of parental MCF-7 cells with the same kinetics (Fig. 2A-red line). As control, we also measured the ability of E2 to increase the number of T47D-1, another ERα expressing breast cancer cell line (Wilson et al., 2004). Figure 2A (purple line) shows that in T47D-1 cells E2 increased the cell number in a time-dependent manner, although with a different kinetics. MCF-7 ERE-NLuc cells were treated with E2 for 24 hours and cell cycle analysis was further performed. E2 augmented the percentage of the cells that incorporated BrdU (Fig. 2B) and increased the number of cells in the S phase of the cell cycle (Fig. 2C) with respect to the control untreated cells. These data demonstrate that E2 induces MCF-7 ERE-NLuc DNA synthesis, cell cycle progression and cell proliferation.

**Figure 2.**
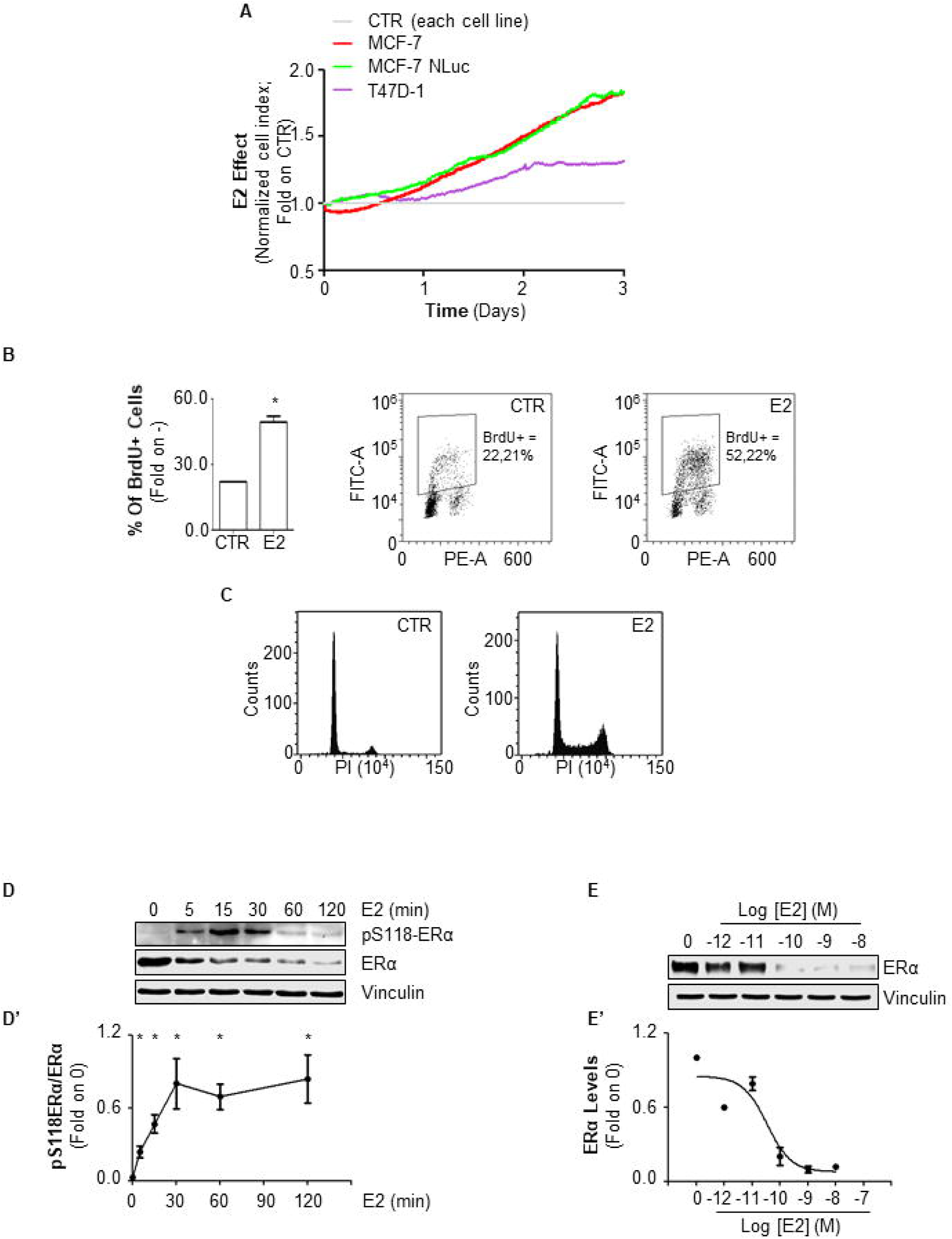
E2 sensitivity of the MCF-7 ERE-NLuc stable cell lines. (A) Growth curves analysis of MCF-7 (red line), MCF-7 ERE-NLuc (green line) and T47D-1 (purple line) cells treated with E2 (10^−9^ M) for 72 hours. Measurement of cell index (C.I.) has been detected every 15 minutes with the xCelligence DP device; for details, please see the method section. Graph shows the E2 effect on C.I. (*i.e*., cell number) calculated for each cell line at each time point with respect to its relative control cells (grey line). The data are the means of two different experiments in which each sample was measured in quadruplicate (for details please see the material and method section). Bromodeoxyurdine (BrdU) incorporation (squares in the plot indicate the BrdU positive events detected by the cytofluorimeter) (B) and cell cycle profile (C) of MCF-7 ERE-NLuc cells treated with E2 (10^−9^ M) for 24 hours. Western blotting (D) and relative densitometric analyses (D’) of S118 phosphorylated (pS118) ERα and ERα expression levels in MCF-7 ERE-NLuc cells treated with E2 at the indicated time points (10^−9^ M); data are the means of three different experiments. Western blotting (E) and relative densitometric analyses (E’) of ERα expression levels in MCF-7 ERE-NLuc cells treated with E2 for 24 hours at the indicated doses; data are the means of three different experiments. The loading control was done by evaluating vinculin expression in the same filter. * indicates significant differences with respect to the CTR or 0 sample. All experiments were performed in triplicates. Data are the mean ± standard deviations with a p value < 0.05.

E2-induced S118 phosphorylation of the ERα is required for receptor transcriptional activity (Ali et al., 1993). Therefore, we tested if E2 could trigger this receptor post-translational modification in MCF-7 ERE-NLuc cells. Time-course analyses revealed that E2 (10^−9^ M) increased the fraction of the S118 phosphorylated ERα in MCF-7 ERE-NLuc cells, which peaked after 30 min of E2 administration and was maintained at least for 120 min (Fig. 2D and D’).

Because E2-induced ERα degradation is intrinsically connected with receptor transcriptional activity (Metivier et al., 2003; Reid et al., 2003) and we noted that E2 also determined a rapid and persistent reduction in ERα intracellular levels (Fig. 2D), we next evaluated the ability of E2 to trigger ERα degradation. Figure 2E and 2E’ show that in 24 hours E2 reduced ERα content in MCF-7 ERE-NLuc cells in a dose-dependent manner with the maximum effect occurring already at 10^−10^ M of E2 administration.

Next, we finally tested the ability of E2 to modulate the expression of the ERE-containing genes pS2/TFF and CatD. As shown in figure 3B and 3B’, the levels of both pS2/TFF and CatD were increased by 24 hours of E2 treatment in MCF-7 ERE-NLuc cells in a dose-dependent manner, with the maximum effect occurring already at 10^−10^ M of E2 administration.

**Figure 3.**
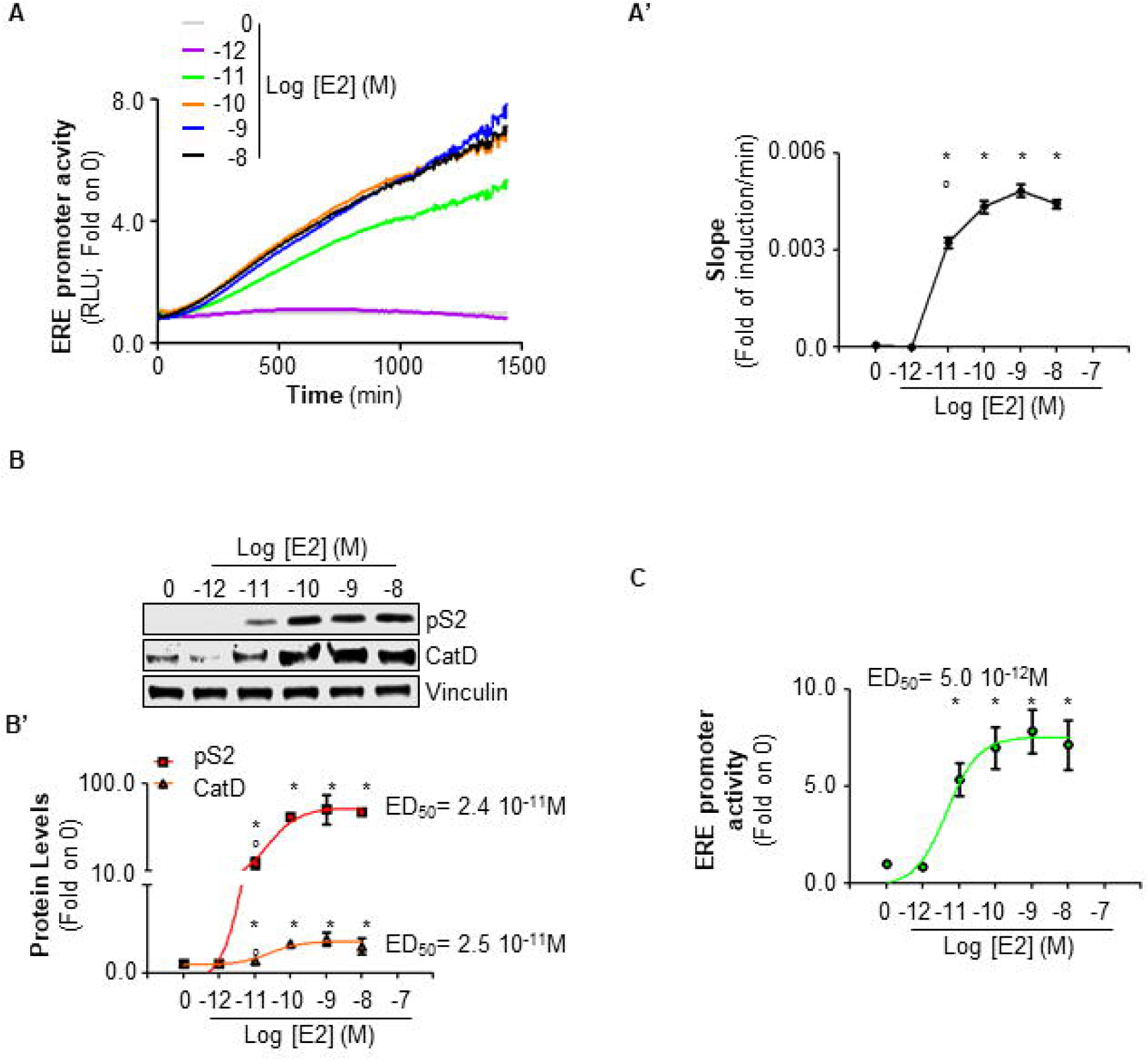
Kinetic analysis of E2 effect in MCF-7 ERE-NLuc stable cell lines. (A) Profile and (A’) relative linear regression (Slope) of ERE-NLuc activity detected in MCF-7 ERE-NLuc cells treated with the indicated doses of E2 in the presence of the live-cell substrate Nano-Glo® Endurazine™. Released light units (RLU) was measured for 24 hours every other 5 minutes in a 37°C and 5% CO_2_ controlled atmosphere. Graph shows the E2 effect calculated at each time point with respect to its relative control sample. The data are the means of three different experiments in which each sample was measured in triplicate (for details please see the material and method section). Western blotting (B) and relative densitometric analyses (B’) of pS2/TFF (red line) and cathepsin D (CatD) (orange line) expression levels in MCF-7 ERE-NLuc cells treated with E2 for 24 hours at the indicated doses; data are the means of three different experiments. The loading control was done by evaluating vinculin expression in the same filter. (C, green line) ERE-NLuc activity detected in MCF-7 ERE-NLuc cells treated with the indicated doses of E2 for 24 hours. * indicates significant differences with respect to the 0 sample; ° indicates significant differences with respect to the 10^−10^ – 10^−8^ M E2-treated samples. Data are the mean ± standard deviations with a p value < 0.01.

Overall, these data indicate that the MCF-7 ERE-NLuc cells respond to E2 as expected for an E2 sensitive cell line derived from the parental MCF-7 cells.

### Real-time measurement of E2-induced ERα transcriptional activity in living cells

To measure the E2-induced ERα transcriptional activity in real-time and in living cells, MCF-7 ERE-NLuc cells were loaded with the live-cell substrate Nano-Glo® Endurazine™ (Hall et al., 2012) in the presence or in the absence of different doses of E2 (from 10^−12^ to 10^−8^ M) and ERE-NLuc-dependent activity (*i.e*., released light units - RLU) was measured for 24 hours every other 5 minutes in a 37°C and 5% CO_2_ controlled atmosphere (for details, please see material and methods section).

As shown in figure 3A, E2 induced a time-dependent increase in ERE-NLuc activity at all the tested doses with a maximal effect occurring already at 10^−10^ M. Notably, cell treatment with 10^−12^ M E2 was ineffective in inducing ERE-NLuc activity (Fig. 3A – purple line). On the contrary, E2-induced activation of ERE-NLuc activity occurred with the same kinetic profile but it reached a lower level of induction when 10^−11^ M E2 was administered to MCF-7 ERE-NLuc cells (Fig. 3A – green line). To evaluate if such differences could be ascribed to a different rate of E2-induced effect, we calculated the slope of the curves (*i.e*., linear regression) obtained by cells treated with E2 in our real-time time course analyses. Figure 3A’ shows a dose-dependent trend in the slope of the curves relative to E2 administration. In particular, no differences were observed among the slopes of the curves that refers to the cells treated with doses of E2 ranging from 10^−10^ to 10^−8^ M while the slope extracted from the 10^−11^ M E2-treated cells was significantly lower (Fig. 3A’). Notably, the slope of the cells treated with 10^−12^ M E2 was identical to that of the control samples (Fig. 3A’).

Next, in order to directly understand if the E2 ERE-NLuc activity correlates with the E2-induced ERE-containing gene expression, we compared the E2 dose-dependent effect on the measured ERE-NLuc activity at the single time point 24 hours (Fig. 3C – green line) with the dose-dependent increase in pS2/TFF and CatD detected in MCF-7 ERE-NLuc cells after 24 hours of E2 administration (Fig. 3B and 3B’). The E2-dependent increase in both ERE-NLuc activity, pS2/TFF and CatD expression was described by a sigmoidal curve typical of the E2 effect. Remarkably, the excitatory dose 50 (ED_50_) calculated for ERE-NLuc activity was lower (ED_50_= 5.0 10^−12^M – 5 pM) than the one calculated for Western blotting analysis of pS2/TFF and CatD cellular levels (ED_50_= 2.5 10^−11^M – 25 pM) (Fig. 3C). Therefore, the ERE-NLuc activity assay is sensitive to E2 administration.

Additionally, we evaluated the involvement of ERα in the E2-induced ERE-NLuc activity. To this purpose we administered different doses (*i.e*., from 10^−7^ to 10^−5^ M) of the ERα antagonist 4OH-tamoxifen (Tam) in the absence or in the presence of E2 (10^−8^ M). As shown in figure 4A, 4B and 4C, the time-dependent linear E2 induction of ERE-NLuc activity was prevented in a dose-dependent manner by Tam administration. Interestingly, different doses of Tam determined different kinetic profiles in MCF-7 ERE-NLuc cells both in the presence and in the absence of E2. While treatment with Tam at 10^−5^ M completely prevented ERE-NLuc activity irrespective of E2 administration (Fig. 4C), lower doses of Tam (*i.e*., 10^−7^ and 10^−6^ M) showed a biphasic kinetic profile both in the presence and in the absence of E2.

**Figure 4.**
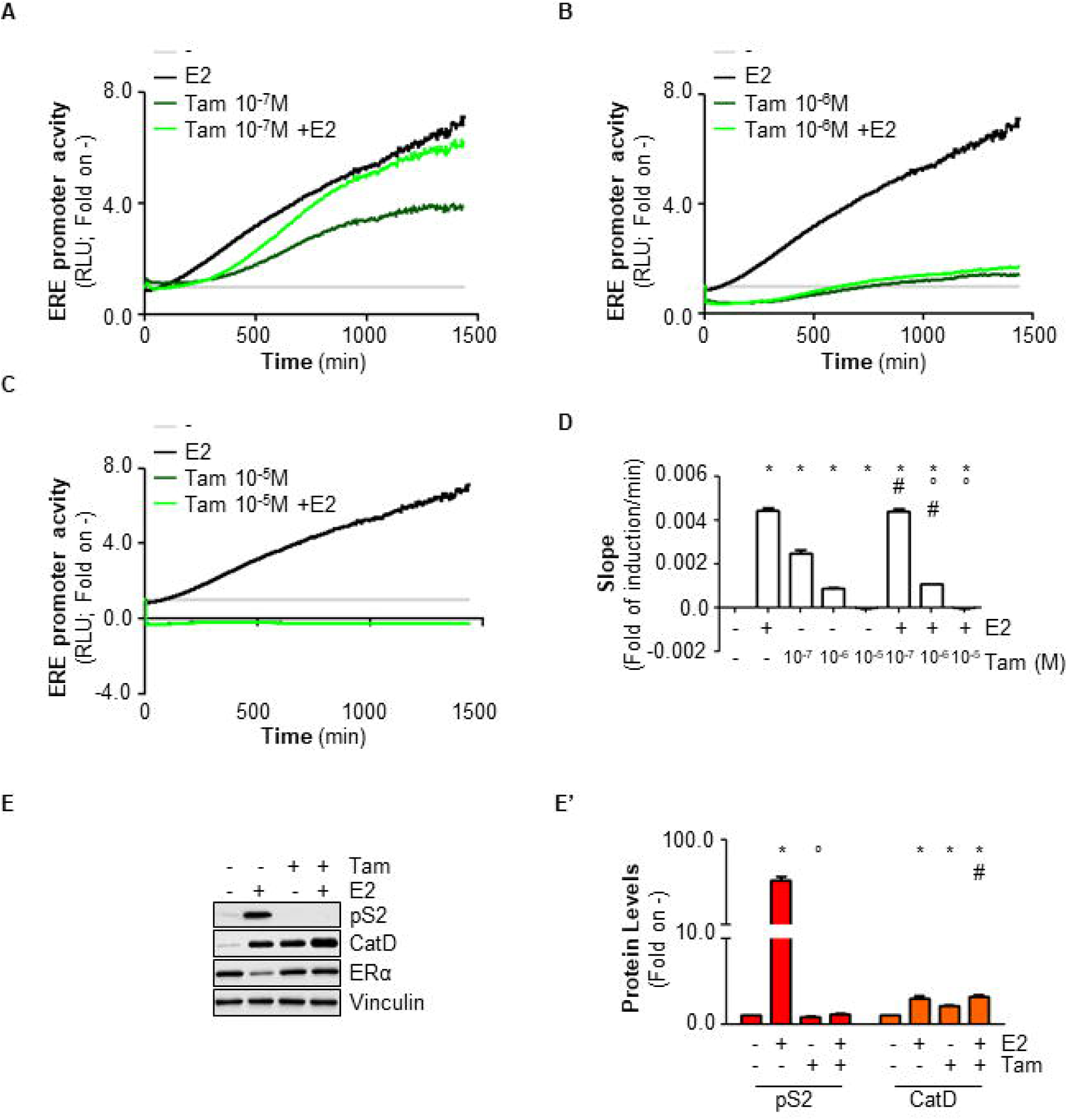
The role of 4OH-tamoxifen on the kinetic analysis of E2 effect in MCF-7 ERE-NLuc stable cell lines. Profile of ERE-NLuc activity detected in MCF-7 ERE-NLuc cells treated with E2 (10^−8^ M) and the live-cell substrate Nano-Glo® Endurazine™ in the presence or in the absence of different doses of 4OH-tamoxifen (Tam): Tam 10^−7^ M (panel A), Tam 10^−6^ M (panel B) and Tam 10^−5^ M (panel C). (D) Linear regression (Slope) relative to real-time measurement of ERE-NLuc activity as depicted in panel A, B and C. Released light units (RLU) was measured for 24 hours every other 5 minutes in a 37°C and 5% CO_2_ controlled atmosphere. Graph shows the E2 effect calculated at each time point with respect to its relative control sample. The data are the means of three different experiments in which each sample was measured in triplicate (for details please see the material and method section). Western blotting (E) and relative densitometric analyses (E’) of pS2/TFF (red bars), cathepsin D (CatD) (orange bars) and ERα expression levels in MCF-7 ERE-NLuc cells treated with E2 (10^−8^ M) in the presence or in the absence of 4OH-tamoxifen (Tam - 10^−7^ M) for 24 hours; data are the means of three different experiments. The loading control was done by evaluating vinculin expression in the same filter. * indicates significant differences with respect to the – sample; ° indicates significant differences with respect to the E2-treated samples. # indicates significant differences with respect to the Tam-treated samples. Data are the mean ± standard deviations with a p value < 0.01 for panel (D) and p < 0.05 for panel (E’).

In a first phase (*i.e*., up to 650 min about 10.8 hours), the values for Tam+E2 treated samples were significantly lower (for Tam 10^−7^ M samples) than those of E2 treated samples or below the control values (for Tam 10^−6^ M samples). For Tam-alone treated samples, instead, we obtained a different behavior as a function of the dose. Indeed, while for Tam 10^−7^ M samples we observed an increase in ERE-NLuc activity, Tam 10^−6^ M-treated samples were significantly below the control levels (Fig. 4A and 4B, respectively).

In a second phase, Tam administration triggered an increase in ERE-NLuc activity up to 1440 min (*i.e*., 24 hours) both in the presence and in the absence of E2 (Fig. 4A and Fig. 4B). Accordingly, the slope of the Tam curves obtained in MCF-7 ERE-NLuc cells were significantly lower than the E2 one but also significantly increased with respect to the control samples (Fig. 4D).

To understand if this Tam-dependent behavior was due to an artifactual measurement of ERE-NLuc activity in our stable MCF-7 cell lines, we measured the E2 effect on pS2/TFF and CatD levels in the presence of Tam (10^−7^ M) in MCF-7 ERE-NLuc cells. As shown in figure 4E and 4E’, the E2-dependent increase in both ERE-NLuc activity, pS2/TFF and CatD expression was prevented by Tam administration (10^−7^ M). However, Tam alone increased the basal levels of CatD (Fig. 4E and 4E’). Notably, a similar stimulatory effect of Tam on ERE-containing genes was already scored in different cell lines (Arao et al., 2011). As expected (Busonero et al., 2019; Leclercq et al., 2006), the E2-induced degradation of ERα was prevented by Tam (Fig. 4E). Altogether, these results show that the E2-induced ERE-NLuc activation depends on ERα transcriptional activity.

Overall, these data demonstrate that E2 induces a rapid and persistent liner increase in ERα transcriptional activity which can be detected at low doses of E2 (*i.e*., between 10^−12^ to 10^−11^ M) and can be prevented by antiestrogens (*e.g*., Tam) and further suggest that the differences in the effects elicited by E2 at different doses could depend on the rate at which E2 induces ERα transcriptional activation.

### Real-time measurement of E2-independent ERα transcriptional activity

Next, we evaluated the ligand-independent ERα transcriptional activation in MCF-7 ERE-NLuc cells. To this purpose we treated cells with epidermal growth factor (EGF), which is known to induce receptor gene transcription in the absence of E2 (Ascenzi et al., 2006).

Because EGF-dependent ERα activation is weak and could be detected only under ERα overexpression conditions (Berno et al., 2008; El-Tanani and Green, 1997), we included a negative control in our experiments by also administering E2 at 10^−13^ M. As expected, E2 induced a time-dependent linear increase in ERE-NLuc activity when cells were treated at hormone dose of 10^−8^ M (black line) but not when 10^−13^ M E2 (orange line) was administered (Fig. 5A). Interestingly, EGF treatment also induced a time-dependent stimulation of ERE-NLuc activity (Fig. 5A). Analysis of both hormone effects on the measured ERE-NLuc activity at the time point 24 hours (Fig. 5B) and the slope of hormone-triggered ERE-NLuc activity curves (Fig. 5C) further confirmed such observations.

**Figure 5.**
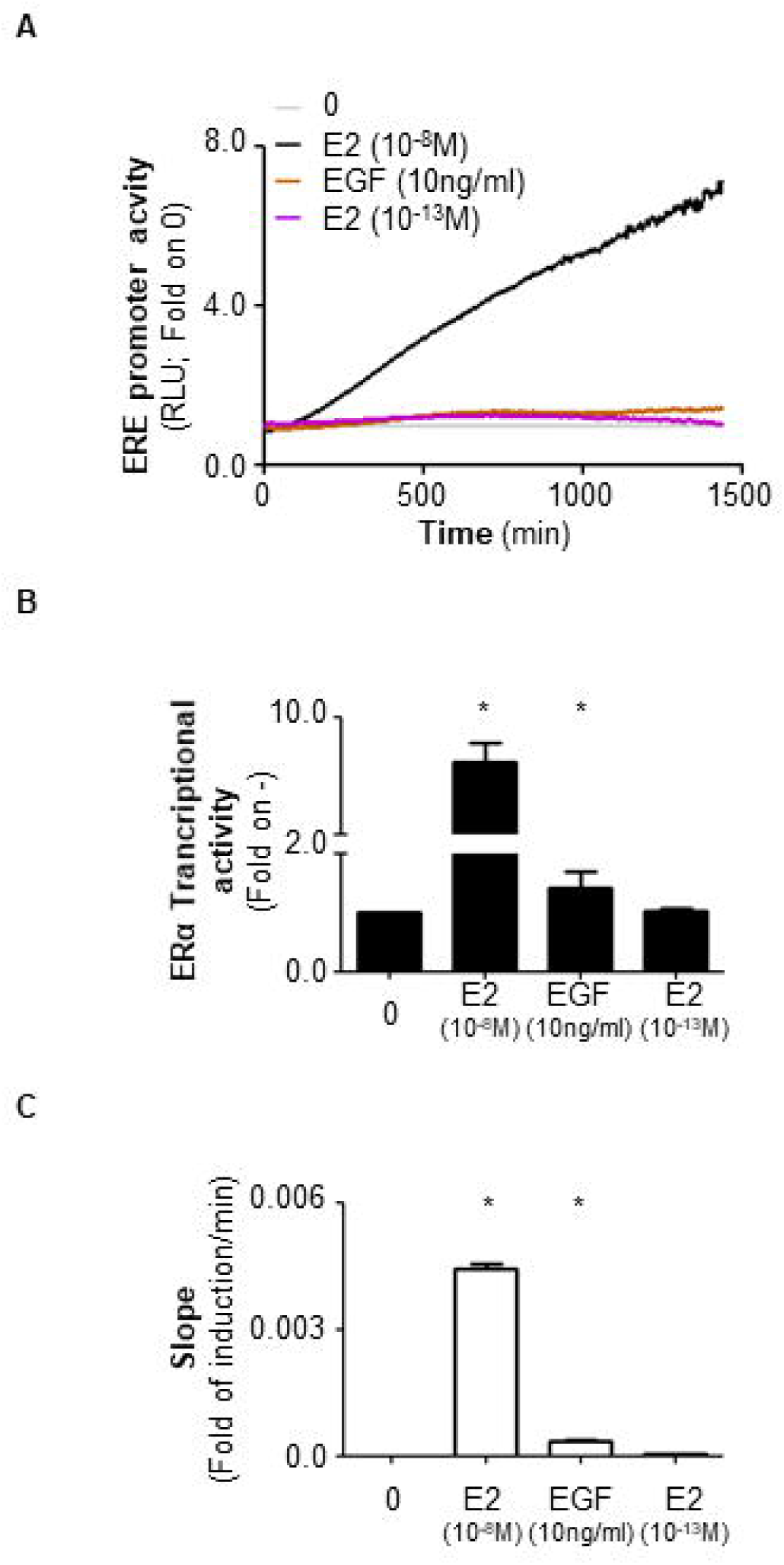
The kinetic analysis of EGF effect in MCF-7 ERE-NLuc stable cell lines. (A) Profile and (C) relative linear regression (Slope) of ERE-NLuc activity detected in MCF-7 ERE-NLuc cells treated with EGF or E2 at the indicated doses and the live-cell substrate Nano-Glo® Endurazine™. Released light units (RLU) was measured for 24 hours every other 5 minutes in a 37°C and 5% CO_2_ controlled atmosphere. Graph shows the E2 effect calculated at each time point with respect to its relative control sample. The data are the means of three different experiments in which each sample was measured in triplicate (for details please see the material and method section). (B) ERE-NLuc activity detected in MCF-7 ERE-NLuc cells treated with EGF or E2 at the indicated doses for 24 hours. * indicates significant differences with respect to the 0 sample. Data are the mean ± standard deviations with a p value < 0.01.

Therefore, in MCF-7 ERE-NLuc cells EGF triggers the activation of the ERα transcriptional function.

### Real-time measurement of the impact of ERα plasma membrane localization on E2-induced ERα transcriptional activity

Plasma membrane localization of the ERα occurs because the receptor is palmitoylated by specific palmityl-acyl-transferases (PATs) on the C residue 447. Inhibition of ERα palmitoylation by either the PAT inhibitor 2-bromo-palmitate (2-Br) or the ERα C447 to alanine (A) mutant (C447A) prevents receptor plasma membrane localization and E2 signaling including ERα transcriptional functions (Acconcia et al., 2005a; Adlanmerini et al., 2014; La Rosa et al., 2012; Pedram et al., 2012; Pedram et al., 2007; Sosa et al., 2019). Therefore, we analyzed the effect of E2 in MCF-7 ERE-NLuc cells in the presence of the PAT inhibitor 2-Br and further evaluated the E2-triggered ERE-NLuc activity of the C447A mutant in transfected HeLa ERE-NLuc cells. Because E2-induced ERα S118 phosphorylation is also required for full receptor transcriptional activity (Ali et al., 1993), we also tested the effects of E2 in ERα S118A mutant transfected HeLa ERE-NLuc cells.

As shown in figure 6A and 6B, 2-Br treatment of MCF-7 ERE-NLuc cells prevented the E2-induced effect on the ERE-NLuc activity and strongly reduced the slope of the curve derived by E2 administration. Similar results were also obtained in wild type (wt) and C447A ERα mutant transfected HeLa ERE-NLuc cells treated with E2 (Fig. 6D and 6E). Notably, as expected for a receptor defective in S118 phosphorylation (Ali et al., 1993), the E2 kinetic profile as well as its relative slope was reduced in S118A ERα mutant transfected HeLa ERE-NLuc cells with respect to HeLa ERE-NLuc cells expressing the wt receptor (Fig. 6D and 6E).

**Figure 6.**
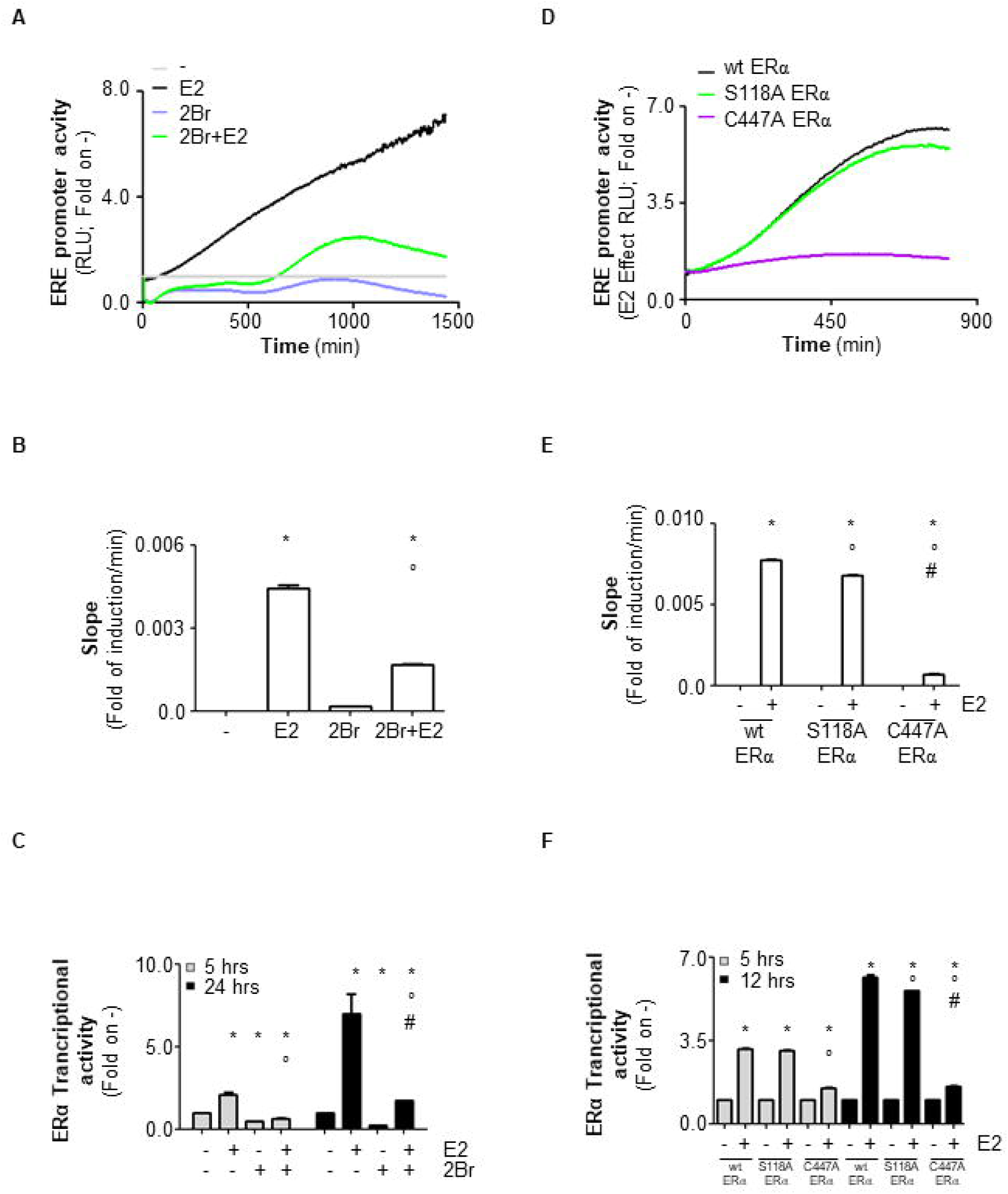
The role of ERα palmitoylation on the kinetic analysis of E2 effect in MCF-7 and HeLa ERE-NLuc stable cell lines. Profile and relative linear regression (Slope) of ERE-NLuc activity detected in the presence of the live-cell substrate Nano-Glo® Endurazine™ either MCF-7 ERE-NLuc cells treated with E2 (10^−8^ M) in the presence or in the absence of 2-bromopalmitate (2-Br - 10^−5^ M) (A, B) or in HeLa ERE-NLuc cells transfected with the pcDNA flag-ERα wild type (wt), the pcDNA flag-ERα C447A and the pcDNA flag-ERα S118A and treated with E2 (10^−8^ M) (D, E). Released light units (RLU) was measured for 24 hours (for MCF-7 ERE-NLuc) and for 12 hours (for HeLa ERE-NLuc) every other 5 minutes in a 37°C and 5% CO_2_ controlled atmosphere. Graph shows the E2 effect calculated at each time point with respect to its relative control sample. The data are the means of two different experiments in which each sample was measured in triplicate (for details please see the material and method section). (C) ERE-NLuc activity detected in MCF-7 ERE-NLuc cells treated with E2 (10^−8^ M) in the presence or in the absence of 2-bromo-palmitate (2-Br - 10^−5^ M) or (F) in HeLa ERE-NLuc cells transfected with the pcDNA flag-ERα wild type (wt), the pcDNA flag-ERα C447A and the pcDNA flag-ERα S118A and treated with E2 (10^−8^ M) at the indicated time points. * indicates significant differences with respect to the – sample; ° indicates significant differences with respect to the E2-treated samples (for MCF-7 ERE-NLuc cells) or to the E2-treated samples in wt ERα (for HeLa ERE-NLuc cells). # indicates significant differences with respect to the 2Br-treated samples (for MCF-7 ERE-NLuc cells) or to the – samples in C447A ERα mutant samples (for HeLa ERE-NLuc cells). Data are the mean ± standard deviations with a p value < 0.01.

Interestingly, we noticed a complex kinetic profile of 2Br-treated MCF-7 ERE-NLuc cells. 2-Br treatment reduced the ERE-NLuc activity below the control. Indeed, within the first 5 hours of treatment E2 was not able to trigger ERE-NLuc activation while from 5 to 24 hours of E2 administration, the hormone stimulated the ERE-NLuc reporter activity (Fig. 6C). On the contrary, the measured E2-dependent ERE-NLuc activation of the C447A ERα mutant in transfected HeLa ERE-NLuc cells was constantly lower than the one detected in the presence of the wt receptor for the entire duration of the analysis (Fig. 6F).

These results confirm that ERα palmitoylation is required for E2-induced ERα transcriptional activity and further suggest that the E2-dpendent activation of ERα plasma membrane localized receptor is necessary for both rapid and persistent activation of ERα transcriptional functions.

### Prediction of dose- and time-dependent E2 effect on ERα transcriptional activity

Finally, based on the slope of the transcriptional profile extracted by our E2 dose-dependent analyses in MCF-7 ERE-NLuc cells (Fig. 2A and 2B), we reasoned that the results obtained from our stable cell lines could allow to predict the time point at which different doses of E2 elicit the same ERE-NLuc activity. In turn, we first calculated the time at which each dose of E2 determines a specific amount of E2 effect (Fig. 7A). On this basis, we hypothesized that 24 hours E2 administration at 10^−11^ M would determine a transcriptional effect equal to those elicited by 18 hours E2 administration at concentration ranging from 10^−10^ M at 10^−8^ M, as detected in ERE-NLuc assays (Fig. 7B).

**Figure 7.**
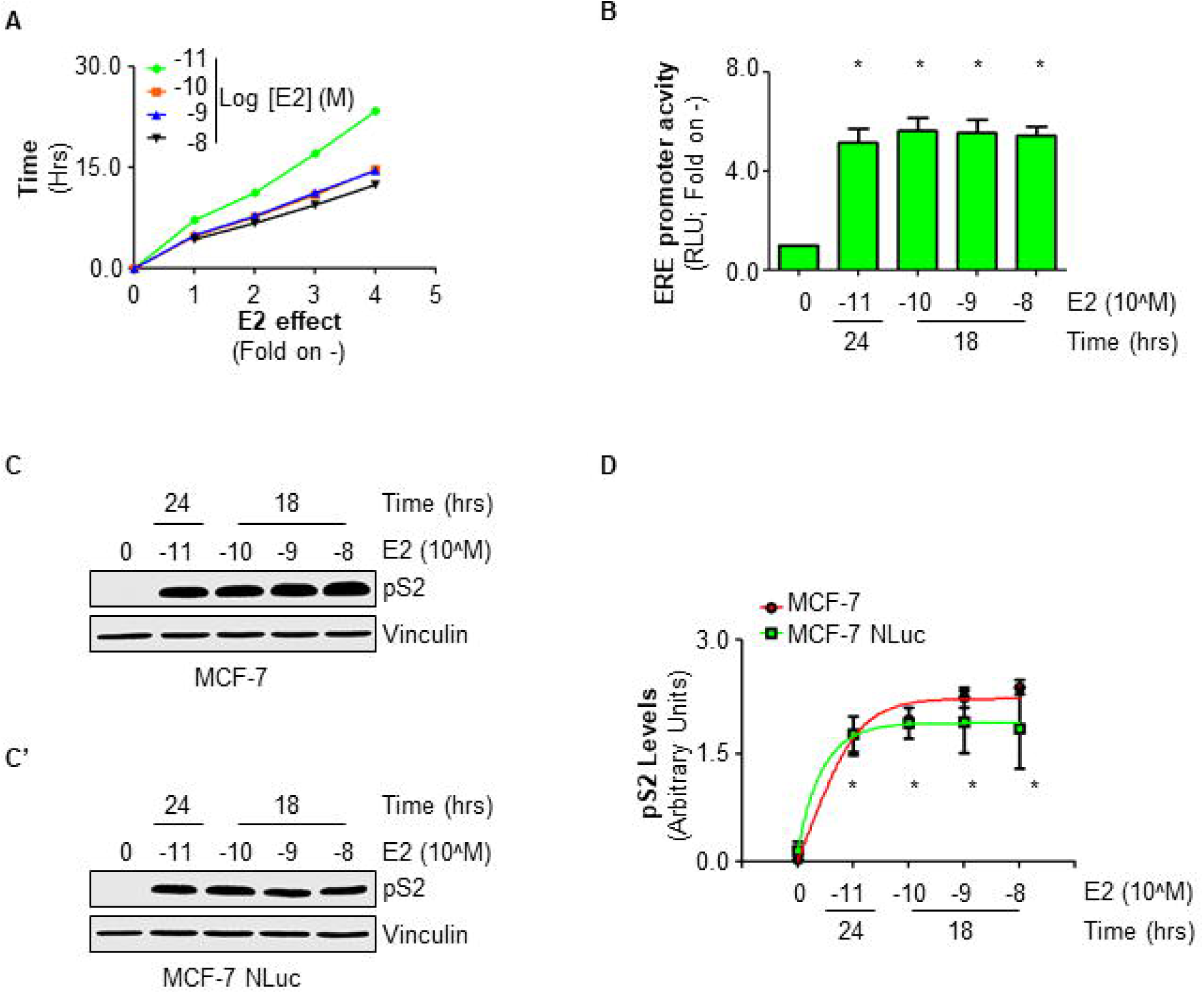
The role of E2 doses and time of E2 administration on ERα transcriptional activity in MCF-7 cells. (A) Time required for E2 dose-dependent induction of ERE-NLuc activity in MCF-7 ERE-NLuc cells. (B) ERE-NLuc activity detected in MCF-7 ERE-NLuc cells treated with E2 at the indicated doses for the indicated times. Western blotting (C, C’) and relative densitometric analyses (D) of pS2/TFF expression levels in MCF-7 (C, red line in D) and in MCF-7 ERE-NLuc cells (C’, green line in D) treated with E2 at the indicated doses for the indicated times; data are the means of three different experiments. The loading control was done by evaluating vinculin expression in the same filter. * indicates significant differences with respect to the – sample. All experiments were performed in triplicates. Data are the mean ± standard deviations with a p value < 0.01 for panel (B) and p < 0.05 for panel (D).

To verify this prediction, we treated both parental MCF-7 and MCF-7 ERE-NLuc cells with E2 at 10^−11^ M for 24 hours and with E2 at both 10^−10^ M, 10^−9^ M and 10^−8^ M for 18 hours and measured the pS2/TFF expression. As shown in figure 7C and 7C’, E2 induced the same accumulation of pS2/TFF intracellular levels at all the tested doses in both parental and artificial MCF-7 cell lines. Remarkably, under these conditions E2 lost its dose-dependent effect (Fig. 3C) on the induction of pS2/TFF expression (Fig. 7D).

These data demonstrate that prolonged treatments of MCF-7 cells with low doses of E2 induce an increase in ERα transcriptional activity identical to that elicited by E2 administered to cells at high concentrations, thus suggesting a critical role for time-dependent effects rather than concentration-dependent effects in the regulation of E2:ERα transcriptional activity.

## Discussion

The main aim of the present work was to identify a method allowing the measurement of ERα transcriptional activity both in real-time and living cells. Here, we constructed a novel plasmid (*i.e*., pGL2Basic Neo_NLucPest_3xERE TATA) containing neomycin resistance cassette and a nanoluciferase-PEST (NLuc) gene under the control of 3 repetition of the classical estrogen response element (ERE) (Fig. 1). The NLuc gene encodes for a modified luciferase gene engineered to be fused in frame with a PEST sequence. This fusion protein returns a high brightness than the classic luciferases and has a shorter protein half-life because of the PEST sequence. Additionally, the substrate for NLuc protein is cell permeable and non-toxic. These biochemical features render such reporter protein particularly suitable for transcriptional studies in living cells (Hall et al., 2012).

In turn, we generated MCF-7 and HeLa cells in which our novel plasmid was stably transfected. MCF-7 cells are ERα expressing cells and are considered standard ‘‘workhorses’’ to study E2-dependent ERα effects (Kao et al., 2009). On the other hand, HeLa cells do not express ERα and therefore do not respond to E2 unless the receptor is exogenously introduced by transient transfection (Acconcia et al., 2005b). The resulting MCF-7-ERE NLuc and HeLa-ERE NLuc stable cell lines are responsive tools to study real-time ERα transcriptional dynamics in living cells under different experimental conditions.

Initial characterization of the stable MCF-7-ERE NLuc cells revealed that E2 induces the proliferation of MCF-7-ERE NLuc cells with a kinetic identical to the one elicited by E2 in the parental MCF-7 cells. Moreover, either MCF-7 cell lines are more sensitive to E2 than the T47D-1 cells, possibly because an higher level of ERα is present in MCF-7 cells than in T47D-1 cells (Fiocchetti et al., 2018; Kao et al., 2009). As expected for an MCF-7-derived cell lines (Dutertre and Smith, 2003; La Rosa et al., 2012; Lannigan, 2003; Leone et al., 2018; Weitsman et al., 2006), we found that the MCF-7-ERE NLuc cells respond to E2 by inducing DNA synthesis, cell cycle progression, the phosphorylation of the ERα on the S118 residue, the degradation of the ERα as well as the increase in the levels of both pS2/TFF and CatD, two classic ERE-containing E2-target genes (Sun et al., 2005). Therefore, we conclude that this novel cell line responds to E2 as the parental MCF-7 cells both in terms of cell proliferation and in terms of intracellular molecular mechanisms of the E2:ERα complex action.

The performance of the stable MCF-7-ERE NLuc cells was evaluated by measuring different parameters of the ERα transcriptional activity. Real-time evaluation of E2 dose-dependent effect indicated that E2 linearly induces ERα transcriptional activity as a function of time. Moreover, we observed that the rate of E2-dependent induction of ERE-based transcription increase in a dose-dependent manner. The excitatory dose 50 (ED_50_) at a single time point (24 hours) following E2 exposure was approximately 5 pM. Notably, these results are similar to those obtained in T47D-1 cells stably transfected with different standard luciferase reporter genes, where the measured ED_50_ for E2 at 24 hours was 6 pM (Legler et al., 1999) or 3 pM (Wilson et al., 2004). Interestingly, the fact that the ED_50_ for E2 at 24 hours for pS2/TFF and CatD measured by means of Western blotting analysis is 25 pM demonstrates that ERE-based assays are more sensitive to variations in ERα transcriptional activity. Accordingly, although the ligand-independent transcriptional activation of the ERα by EGF is weak and could be detected only in cells overexpressing the receptor (Berno et al., 2008; El-Tanani and Green, 1997), we were able to profile the temporal dynamics of the EGF-induced ERα transcriptional activity also in endogenously expressing ERα cells.

Time-dependent E2-induced ERα transcriptional activity was prevented by Tam in MCF-7-ERE NLuc cells. Indeed, the rate of E2-transcriptional induction was strongly reduced in the presence of Tam. However, analysis of the time-dependent transcriptional profile of Tam-treated MCF-7-ERE NLuc cells identified a biphasic kinetic both in the presence and in the absence of E2. Tam is the prototype selective estrogen receptor modulator (SERM). It works by binding to the ERα and inhibiting its transcriptional activity through receptor structural modifications (Brzozowski et al., 1997). In our real-time live-cell analyses, we observed a first phase in which the ERα transcriptional activity was reduced at all the tested doses. However, at later time points, ERα transcriptional activity was stimulated by Tam although the E2 effect was reduced for low doses of Tam (*i.e*., 10^−7^ M) and completely abolished for high doses of Tam (*i.e*., 10^−6^ and 10^−5^ M). Accordingly, although the E2 effect in inducing the expression of CatD and pS2/TFF was reduced, CatD but not pS2/TFF basal expression levels were increased by Tam (10^−7^ M). Although this discrepancy is difficult to reconcile, it is however possible to speculate that different E2-target genes have different sensitivity to E2 administration because they respond to a different amount of active/inactive ERα (please see below). Indeed, basal Tam-dependent ERα activation could be ascribed to the ability of Tam to induce an accumulation in ERα intracellular levels (Busonero et al., 2019; Leclercq et al., 2006). The increased number of ERα molecules would compensate for the inhibitory effect of Tam but only at later time points. On the contrary, the inhibition of receptor transcriptional activity is immediate and consistent with the ability of Tam to bind ERα and to induce an antagonist receptor conformation in a time-frame compatible with those of ligand:receptor association (Brzozowski et al., 1997). This potential biochemical mechanism appears to be supported by the fact that high doses of Tam (*i.e*., 10^−5^ M) do not show a biphasic kinetic profile and completely prevent the basal and E2-induced ERα transcriptional activity. However, the observed stimulatory activity of Tam on ERα transcriptional activity is consistent with already reported observations (Arao et al., 2011).

Real-time analysis of the impact of ERα plasma membrane localization on E2-induced ERα transcriptional activity showed that the rate of this process was strongly dampened in cells in which ERα palmitoylation was prevented. More interestingly, the ERα transcriptional activity showed a biphasic kinetic profile in MCF-7-ERE NLuc cells treated with the PAT inhibitor. Inhibition of ERα palmitoylation results in an inhibition of the E2-induced extra-nuclear signaling which are required for the activation of receptor transcriptional activity and for the completion of many different E2-dependent physiological functions both in cell lines and *in vivo* (Acconcia et al., 2005a; Adlanmerini et al., 2014; La Rosa et al., 2012; Pedram et al., 2012; Pedram et al., 2007; Sosa et al., 2019). Our time-dependent live-cell analyses confirm that the contribution of the rapid signaling pathways originating from the activation of the ERα at the plasma membrane is critical for full receptor transcriptional activity. However, data presented here further demonstrate that besides being a pre-requisite for the rapid initial stages of E2-induced ERα transcriptional activation (La Rosa et al., 2012), the E2:ERα plasma membrane signaling is also absolutely required for the prolonged effect of ERα as a ligand-induced transcription factor. In support to this notion, the time-dependent kinetic profile of E2-induced transcriptional activity of the palmitoylation defective ERα C447A mutant measured in HeLa-ERE NLuc cells was completely flat. Therefore, we conclude that the plasma membrane localization of the ERα is necessary and sufficient for the induction of the receptor transcriptional activity both for rapid and prolonged times of E2 administration.

Finally, we predicted the time-dependent transcriptional behavior of different doses of an ERα ligand. Indeed, we have calculated the time required for different E2 concentrations to reach the same transcriptional effects and effectively reported that treatments with different E2 concentrations at different time points of both MCF-7-ERE NLuc and parental MCF-7 cells produced the same increase in pS2/TFF expression. Physiological blood concentration of E2 fluctuates in healthy pre-menopausal woman between picomolar and nanomolar concentrations. However, the peak of E2 concentration in blood lasts for a maximum time of 48-72 hours. This peak is however sufficient to induce all the physiological uterine modifications (Zittermann et al., 2000). Our results suggest that rather than the chronic exposure to E2, the time of E2 administration is critical to achieve the maximal ERα transcriptional activity. In this respect, the pulsatile nature of E2 action under physiological conditions would support such concept. However, future investigations are required to establish if short E2 treatments could lead to fully and prolonged activation of ERα action.

In conclusion, we report a method to study the real-time kinetics of E2:ERα transcriptional activity in living cells using the generated MCF-7-ERE NLuc and HeLa-ERE NLuc stable cell lines. Here, we only measured the classic parameters of the E2 signaling (*i.e*., ligand-dependent and independent functions; the impact of ERα antagonists and of receptor plasma membrane localization) on ERα transcriptional activity in MCF-7-ERE NLuc or in HeLa-ERE NLuc cells. Nonetheless, our stable cell lines are suitable for diverse analyses. MCF-7-ERE NLuc cells can be easily used for drug discovery by setting up high-throughput screenings for the identification of novel ERα antagonists with specific kinetic profiles. In addition, these cell lines could facilitate the analysis of the transcriptional effects elicited by known or unknown endocrine disruptors or environmental contaminants that bind ERα and have a weak estrogenic activity. In this respect, the use of a real-time live-cell assay could be an advantage in understanding complex dose-dependent effects of such receptor ligands (*e.g*., U-shaped curves) (Acconcia et al., 2016; Acconcia et al., 2015; Marino et al., 2012). MCF-7-ERE NLuc cells are also in principle suitable for the analysis of the physiological involvement of a specific protein on the regulation of E2:ERα transcriptional activity (*e.g*., single or high-throughput screening with siRNA oligonucleotides). On the other hand, HeLa-ERE NLuc cells are transfectable with different kinds of ERα (or ERβ) deletion or point mutants and can be used to connect ERα (or ERβ) structural biochemical determinants with receptor transcriptional activity. Finally, this experimental model could be applied to other nuclear receptors (*e.g*., ERβ, androgen receptor, glucocorticoid receptor) enlarging our knowledge on the dynamic of nuclear receptor transcriptional activity and its modulation by different endogenous or exogenous ligands.

Therefore, these stable cell lines represent new models for the real-time live-cell analysis of the kinetic aspects of E2 signaling under physiological or pathophysiological conditions.

## Data availability statement

Research data not shared. However, the data sets used and/or analyzed during the current study are available from the corresponding author on reasonable request.

Contract grant sponsor: Associazione Italiana Ricerca sul Cancro AIRC, Fondo di finanziamento per le attività base di ricerca (FFABR) to FA. The Grant of Excellence Departments, MIUR (ARTICOLO 1, COMMI 314 – 337 LEGGE 232/2016) to Department of Science.

## Author contribution statement

M.C. performed most of the work (generation of stable cell lines; real-time live cell experiments; most of Western blotting analyses); S.L. performed all cell cycle analyses. S.B. and C.B. generated the expression vectors required for transfection and performed some of Western blotting analyses. M.C., S.L., S.B. and C.B. to figure preparation and manuscript proof-reading. F.A. designed the research and wrote the paper.

## Acknowledgments

The research leading to these results has received funding from AIRC under IG 2018 - ID. 21325 project – P.I. Acconcia Filippo. This study was also supported by grants from Ateneo Roma Tre and Fondo di finanziamento per le attività base di ricerca (FFABR) to FA. The Grant of Excellence Departments, MIUR (ARTICOLO 1, COMMI 314 – 337 LEGGE 232/2016) to Department of Science, University Roma TRE is also gratefully acknowledged. The Authors declare no conflict of interest.

